# Experimental evolution of reproductive isolation from a single natural population

**DOI:** 10.1101/436287

**Authors:** Scott M. Villa, Juan C. Altuna, James S. Ruff, Andrew B. Beach, Lane I. Mulvey, Erik J. Poole, Heidi E. Campbell, Kevin P. Johnson, Michael D. Shapiro, Sarah E. Bush, Dale H. Clayton

## Abstract

Ecological speciation occurs when local adaptation generates reproductive isolation as a by-product of natural selection^1–3^. Although ecological speciation is a fundamental source of diversification, the mechanistic link between natural selection and reproductive isolation remains poorly understood, especially in natural populations^2–6^. Here we show that experimental evolution of parasite body size over four years (ca. 60 generations) leads to reproductive isolation in natural populations of feather lice on birds. When lice are transferred to pigeons of different sizes they rapidly evolve differences in body size that are correlated with host size. These size differences trigger mechanical mating isolation between lice that are locally adapted to the different sized hosts. Size differences among lice also influence the outcome of competition between males for access to females. Thus, body size directly mediates reproductive isolation through its influence on both inter-sexual compatibility and intra-sexual competition. Our results confirm that divergent natural selection acting on a single phenotypic trait can cause reproductive isolation to emerge from a single natural population in real time.

---

Feather lice (Insecta: Phthiraptera: Ischnocera) are host-specific parasites of birds that feed and pass their entire life cycle on the body of the host. Members of the genus *Columbicola* are parasites of pigeons and doves (Columbiformes) that feed on the downy regions of feathers, causing energetic stress and a reduction in host fitness through reduced survival and mating success^7^. Pigeons combat feather lice by removing them with their beaks during regular bouts of preening. *Columbicola columbae*, a parasite of pigeons (*Columba livia*), avoids preening by hiding in spaces between adjacent feather barbs (Fig. 1a); preening selects for *C. columbae* that fit between the barbs^7,8^. In the absence of preening, large bodied lice are favored because of a size-fecundity correlation; all else being equal, large lice lay more eggs than small-bodied lice^9^. These opposing selective forces explain the high correlation between the body sizes of different species of pigeons and their host-specific species of *Columbicola*; this macroevolutionary pattern is known as Harrison’s Rule^7,10^.

**Figure 1.**
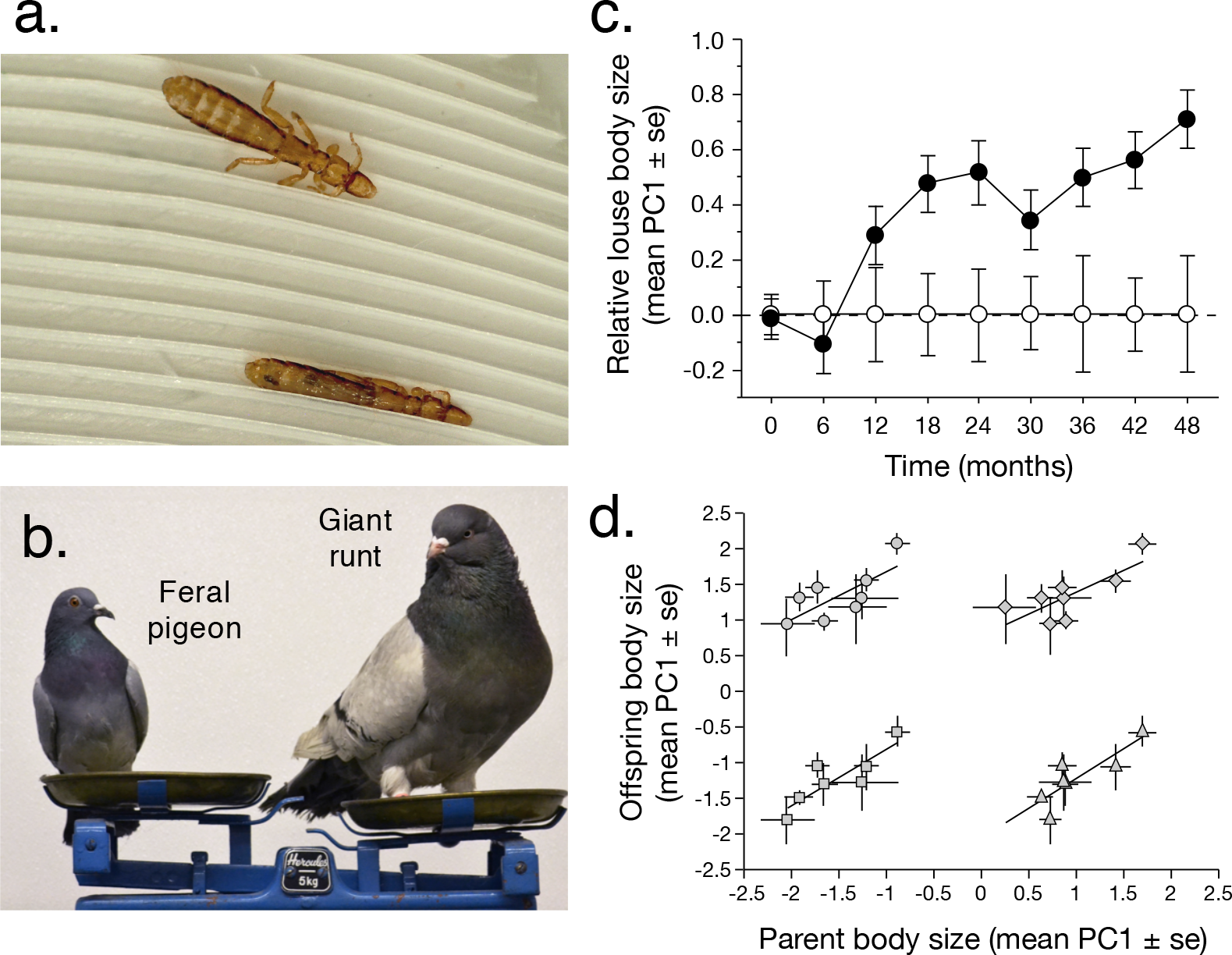
Experimental evolution of *Columbicola columbae* body size. (a) Live *C. columbae* walking on the surface of a feral pigeon wing feather (top), and inserted between adjacent feather barbs (bottom) to escape host preening. (b) Relative sizes of a feral pigeon (ca. 340g) and a domesticated giant runt pigeon (ca. 1100g), both *Columba livia*. (c) Increase in the relative size of *C. columbae* on giant runts (black circles) over four years (ca. 60 louse generations), compared to the size of lice on feral pigeons (white circles, set to zero) (LMM, n = 3096, t = 3.15, *P* = 0.002). (d) Common garden experiment showing that *C. columbae* body size is heritable. Each point compares the mean (± se) body size of parental and offspring cohorts on a single common garden feral pigeon. Parent and offspring size are highly correlated in all cases. Points are: daughters vs. fathers (circles, upper left; linear regression, r = 0.73, df = 7, F = 7.13, *P* = 0.037), daughters vs. mothers (diamonds, upper right; r = 0.77, df = 7, F = 8.65, *P* = 0.026), sons vs. fathers (squares, lower left r = 0.86, df = 6, F = 13.90, *P* = 0.014), and sons vs. mothers (triangles, lower right r = 0.84, df = 6, F = 12.61, *P* = 0.016).

Because feather lice are permanent parasites that pass their entire life cycle on the body of the host, they can be maintained under natural conditions on captive birds^7^. This fact, coupled with their short 24-day generation time, makes feather lice tractable for experimental evolution studies of reproductive isolation. Feather lice essentially provide an ecological intermediate between conventional lab models, such as *Drosophila*^11^, pea aphids^12^, and *Tribolium*^13^, and field-based models, such as Darwin’s finches^14^, *Heliconius* butterflies^15^, *Rhagoletis* flies^16^, and threespine sticklebacks^17^.

To test for adaptation in response to host body size, we conducted a four-year experiment (ca. 60 louse generations) using *C. columbae* placed on captive pigeons of different sizes. We transferred lice from wild caught feral pigeons to giant runts, a domesticated breed of pigeon that is three-fold larger than feral pigeons (Fig. 1b). We also transferred *C. columbae* to feral pigeon controls. We quantified the size of lice by measuring louse body length, metathorax width, and head width. These measures are highly correlated, so we used the first principal component (PC1) as an index of overall louse size (Table S1). Over the course of four years, lice on giant runts increased in size, relative to lice on feral pigeon controls (Fig. 1c, Tables S2-S5). Mid-way through the study (two years), we performed a common garden experiment and confirmed that louse body size has a heritable component (Fig. 1d). In summary, our four-year experimental evolution study shows that host-imposed natural selection drives rapid local adaptation in the body size of lice.

Body size also plays a role in the reproductive biology of *C. columbae*. Male and female *C. columbae* are sexually dimorphic in overall body size (Fig. 2a), and in the structure of the male antennae (Fig. 2b), which are used to grasp the female during copulation (Fig. 2c). Behavioral observations of live *C. columbae* suggest that the extent of body size dimorphism may influence copulation success. During copulation between individuals showing “typical” dimorphism (Table S6), males use their antennae to grasp the female’s metathorax while aligning the tip of their abdomen with the female’s abdomen (Fig. 2c, 3a,b; Video S1). When males are too small or too large, relative to the female, they have difficulty copulating (Fig. 3c-f; Videos S2,S3). These observations suggest that increases in the body size of *C. columbae* on giant runts may reduce their ability to copulate with smaller lice from feral pigeons. Thus, reproductive isolation may evolve as a direct result of adapting to a new environment^3,18,19,20^.

**Figure 2.**
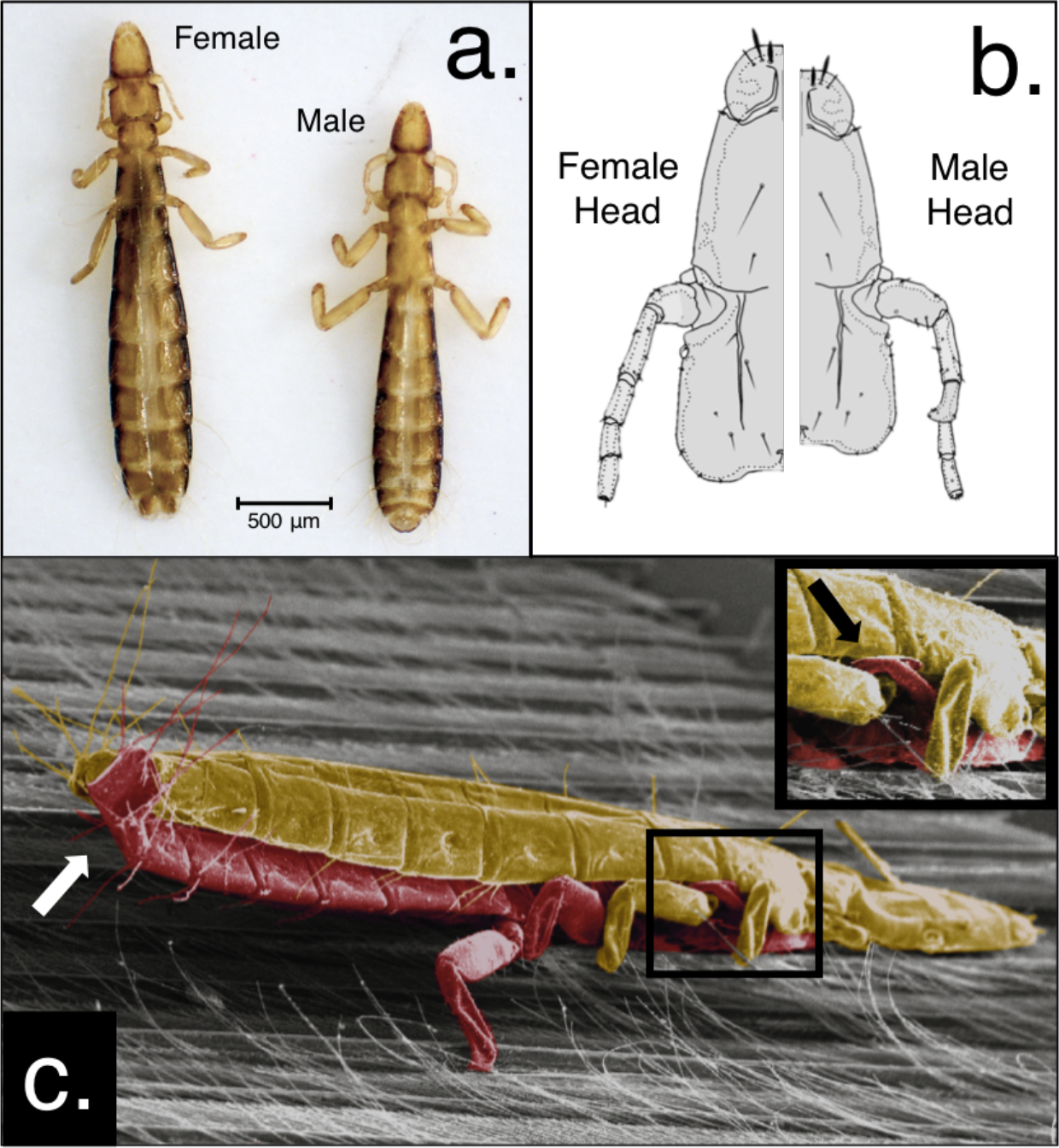
Sexual size dimorphism of *C. columbae*. (a) females are typically 13% larger than males. (b) Sexually dimorphic heads showing male antenna with larger scape (first segment) and inward pointing spur on the third segment. (c) Colorized SEM of *C. columbae* copulating on a pigeon feather: male (red) grabbing the female (gold) with his antennae (black arrow; inset), while curling the tip of his abdomen dorsally to contact the tip of her abdomen (white arrow).

**Figure 3.**
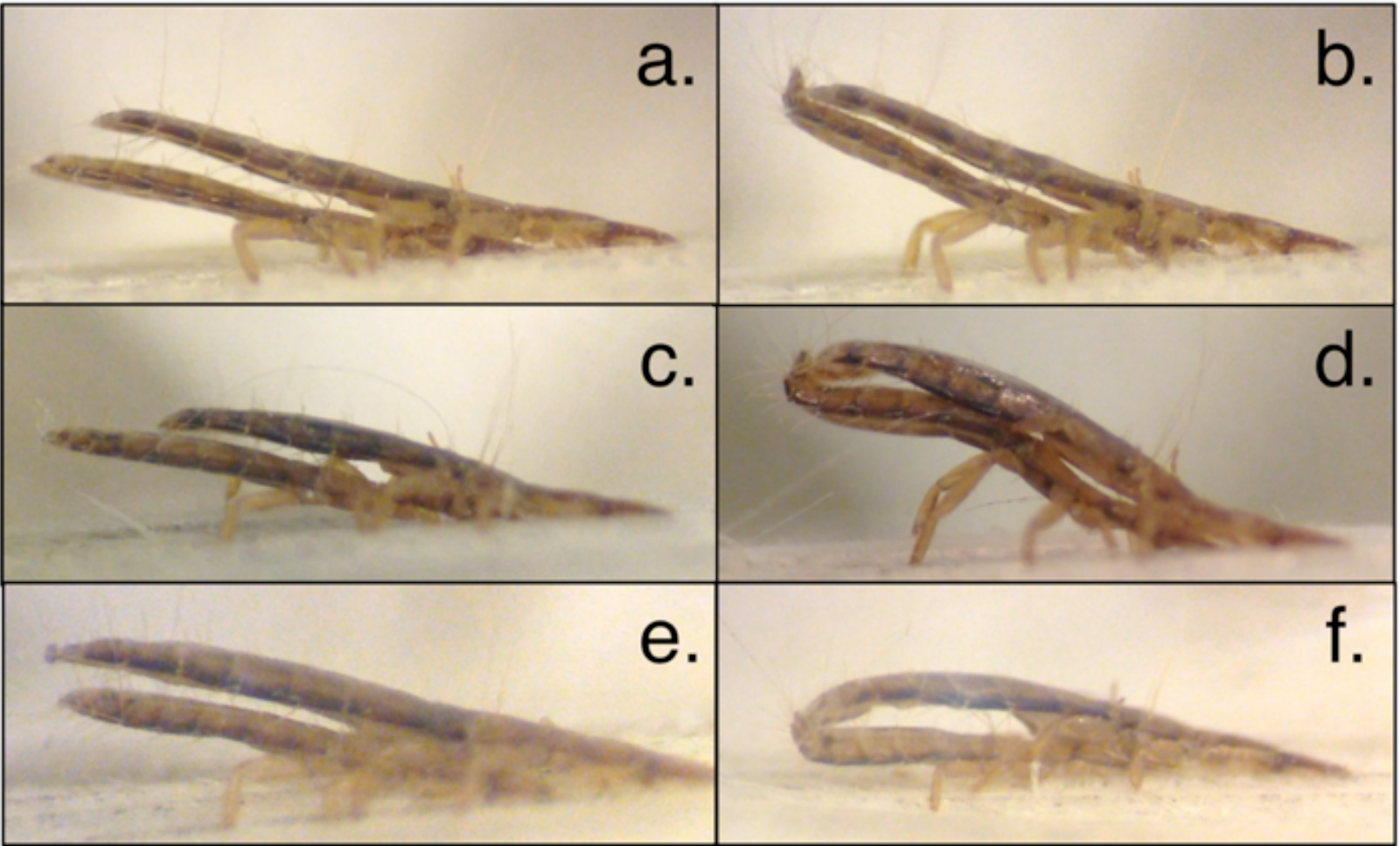
Influence of body size on mating position in *C. columbae* (female on top, male on bottom in all photos). (a-f) Still photographs from videos (see Supplemental Information for videos). (a,b) Abdomens are parallel during copulation in pairs of lice with typical dimorphism (Video S1). (c,d) Relatively large males seldom succeed in copulating, but when they do they are S-shaped (Video S2). (e,f) Females copulating with relatively small males, which is also rare, are arched during copulation (Video S3). Dimorphism scores (male length - female length): a,b = −346μm; c,d = −197μm; e,f = −561μm.

We conducted a series of experiments to test the effect of body size on louse reproductive compatibility (Fig. 4). First, we quantified time spent copulating by pairs of lice that vary in degree of dimorphism. Lice were filmed in mating arenas on detached feathers. Pairs of lice with typical dimorphism (Table S6) copulated for significantly longer than “mismatched” pairs with more or less dimorphism (Fig. 4a). Observations of lice in mating arenas showed that, although virtually all males attempted to copulate, individuals that were too large or too small, relative to females, had difficulty. On average, mismatched pairs spent 70% less time copulating than typical pairs. Thus, copulation time is a function of the relative dimorphism of male and female lice (Fig. 4a). Copulation time predicts reproductive success in insects^21^.

**Figure 4.**
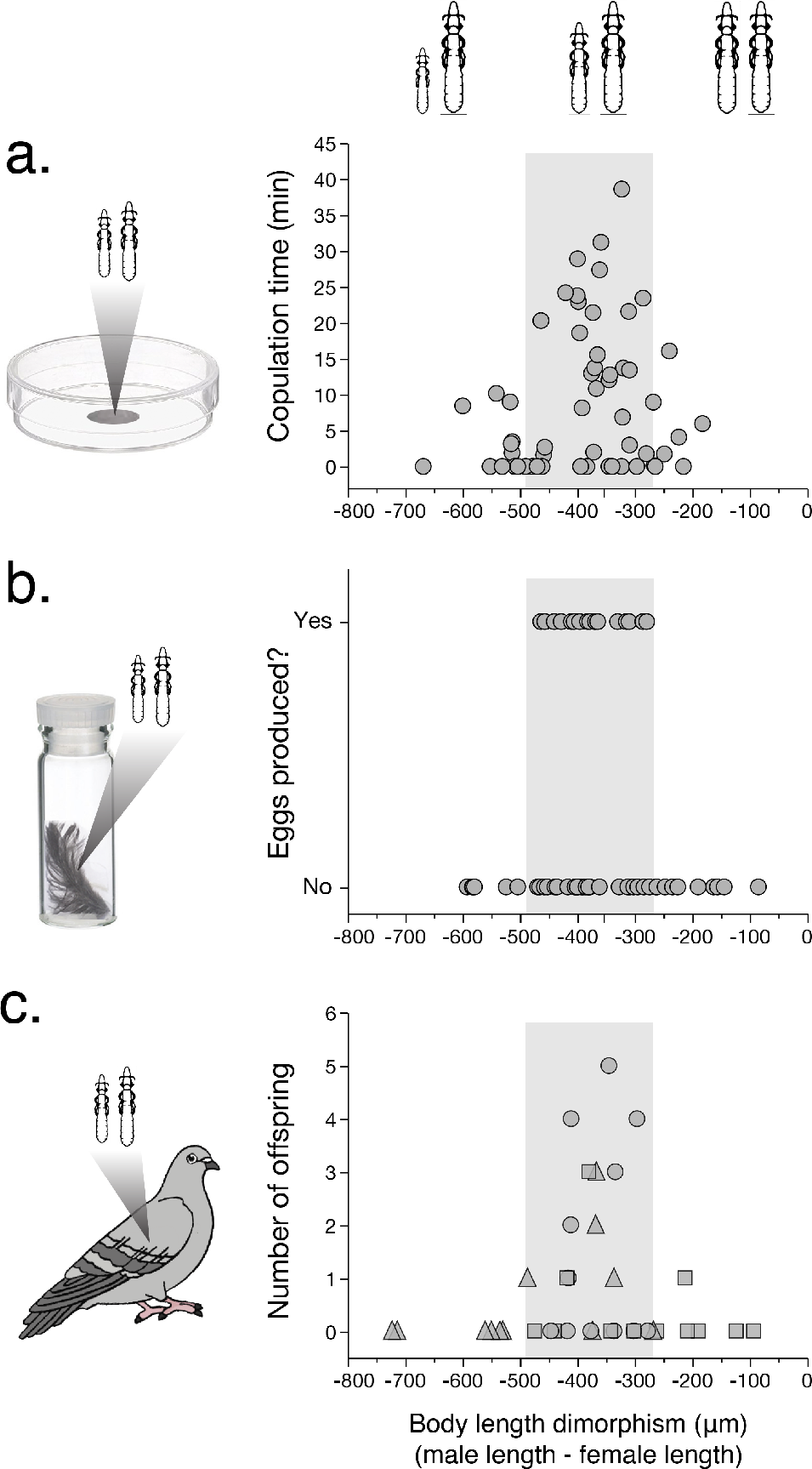
Reproductive performance of *C. columbae* differing in size dimorphism. Values on the *x*-axis indicate the difference in body length of the male relative to that of the female (e.g. “-400 μm” represents a trial in which the male was 400 μm shorter than the female). Grey shaded region in each plot represents the typical dimorphism of lice (Table S6); relatively small males at left, large males at right (illustrations not to scale). (a) Copulation time of lice in mating arenas was correlated with size dimorphism (quadratic regression, n = 56, F = 3.53, *P* = 0.03). (b) Eggs were produced only by pairs with typical dimorphism (ordinal logistic regression, n = 58, Chi Square = 11.20, *P* = 0.003); 17 of 42 typical pairs (40.5%) produced eggs, but none of 16 pairs with relatively small or large males produced eggs. (c) Reproductive success of 36 pairs of lice from feral or giant runt pigeons transferred to 36 louse-free feral pigeons: 12 pairs included a male giant runt louse and a female feral pigeon louse (squares); twelve pairs included a male feral pigeon louse and a female giant runt louse (triangles); twelve (control) pairs included a male feral pigeon louse and a female feral pigeon louse (circles) (Table S7). Twelve of 23 (52.2%) typical pairs produced offspring, whereas just one of 13 pairs (7.6%) with relatively small or large males produced any offspring (Fisher’s exact test, n = 36, *P* = 0.01). Reproductive success was governed by the relative size of the male and female lice, independent of the type of host on which they evolved (cf. circles, triangles, and squares).

We also tested the effect of size dimorphism on the production of eggs by lice. Pairs of virgin lice were placed in vials of feathers kept in an incubator at optimal temperature and humidity^22^. The vials were checked daily for several weeks until the female in each vial died. Pairs of lice with typical dimorphism (Table S6) were significantly more likely to produce eggs than mismatched pairs (Fig. 4b). Nearly all of the eggs (92%) had developing embryos. Thus, egg production is also a function of relative dimorphism.

Next we tested the effect of variable dimorphism on the reproductive success of lice on live pigeons. Single pairs of lice were placed on individual pigeons for two weeks to quantify the number of F1 offspring they produced. Again, typical pairs of lice had significantly more offspring than mismatched pairs (Fig. 4c). Relative size dimorphism thus dictates the number of offspring produced by lice under natural conditions on live birds. The results of the three male-female pair experiments (Fig. 4a-c) show that local adaptation triggers reproductive isolation in cases where male and female lice are either too different, or too similar, in body size.

Sexual size dimorphism also influences intra-sexual competition for mates^23,24,25^. We tested if the degree of size dimorphism influences male-male competition in *C. columbae* by allowing two males to compete for single females in mating arenas. Typical males spent significantly more time copulating than either small or large males (Fig. 5a-c). During these trials we observed males trying to displace other males already *in copula* by wedging themselves between the first male and the female (Video S4). We also observed behaviour consistent with mate guarding by males following copulation. Thus, sexual selection magnifies the consequences of local adaptation for the evolution of reproductive isolation in *C. columbae* lice^6,25,26,27^.

**Figure 5.**
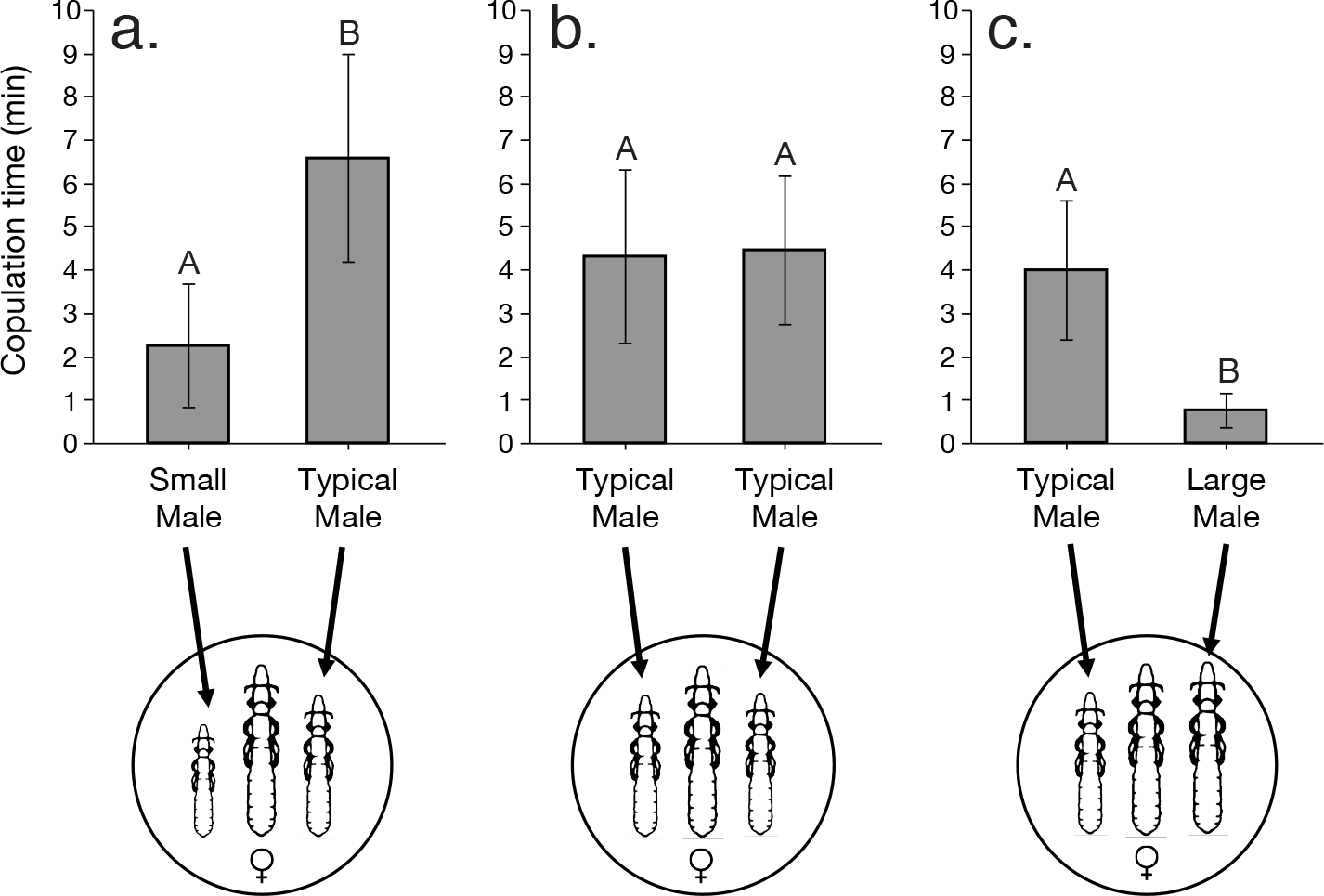
Relationship of size dimorphism to male-male competition in *C. columbae* (lice illustrations not to scale). When males of different sizes were combined with a single female in mating arenas, typical sized males spent more time copulating than relatively small (a) or large (c) males; Wilcoxon Signed Rank Tests: n = 10, S = 24.50, *P* = 0.02 for (a), and n = 10, S = 22.50, *P* = 0.03 for (c). When males of the same (typical) size were combined with a single female (b), the males did not differ significantly in copulation time; n = 10, S = 13.00, *P* = 0.25. Different letters indicate significant differences for *P* < 0.05.

Body size governs the survival of lice on different sized hosts, as well as the reproductive compatibility of lice locally adapted to those hosts. Thus, body size in lice is consistent with a “magic trait” model of ecological speciation^3,20,28^. Experimental divergence in size over four years (ca. 60 generations) led to partial reproductive isolation between populations adapted to large and small-bodied hosts. The rate of change we observed, together with projected increases in size^7,10^, suggest that lice on giant runts could be completely isolated from lice on feral pigeons within 450 generations (30 years). This prediction assumes sustained selection on body size, and the maintenance of additive genetic variation in body size. Our results show that the early stages of ecological speciation can be fast, taking place on the same time scale as adaptive divergence^3,27^.

The rapid evolution of body size and the emergence of reproductive isolation between lice on different sized hosts presumably resembles the consequences of host switches by lice in nature. Feather lice are relatively host specific and have phylogenies that are congruent with those of the host, owing to repeated bouts of host-parasite cospeciation^7^. Nevertheless, lice do sometimes switch host lineages^7^. If lice switch to a new host that differs in body size from the original host, then local adaptation should lead to rapid evolution of louse body size. This, in turn, should lead to the emergence of reproductive isolation between populations of lice that come back into secondary contact. If the change in body size is large enough over time, then complete prezygotic reproductive isolation could result in the formation of a new species. However, even if the difference in body size does not yield complete reproductive isolation, the interruption in gene flow created by partial mechanical isolation could facilitate the evolution of post-zygotic barriers over longer periods of time^1,26,27^.

Our study emphasizes the benefit of examining ecological speciation in systems during the earliest stages of divergence. Most studies of ecological speciation retrospectively investigate closely related taxa that have already diversified^1,4,11–17,26,27^. We took a complementary approach and experimentally triggered diversification in the descendants of a single population living under natural conditions in real time. This approach allowed us to identify the specific trait that links adaptation to reproductive isolation. By showing that local adaptation leads directly to the rapid emergence of reproductive isolation, our results confirm a fundamental prediction of ecological speciation theory^1,28,29,30^.

## METHODS

Detailed methods are provided at the end of the manuscript.

## SUPPLEMENTAL INFORMATION

Supplemental information includes seven tables at the end of this manuscript. Supplemental information also includes four linked videos. Legends for each video are at the end of this manuscript.

## ACKNOWLEDGEMENTS

We thank F. Adler, D. Benevides, M. Evans, G. Goodman, S. McNew, N. Phadnis, J. Seger, and E. Waight for discussion and other assistance. All procedures followed guidelines of the Institutional Animal Care and Use Committee of the University of Utah. This work was supported by National Science Foundation DEB-1342600.

## AUTHOR CONTRIBUTIONS

S.M.V, S.E.B, K.P.J, M.D.S, and D.H.C. designed the experiments. S.M.V., J.C.A., A.B.B, L.I.M., E.J.P., and H.E.C. collected the experimental data. S.M.V., J.S.R., S.E.B, and D.H.C. analyzed the data. S.M.V., J.S.R., K.P.J, M.D.S., S.E.B. and D.H.C. contributed to writing of the manuscript. J.C.A., A.B.B, L.I.M, E.J.P., and H.E.C took and analyzed the digital photographs of lice used in experiments. S.M.V., S.E.B, and D.H.C. prepared figures.

## METHODS

### Elimination of “background” lice

Before using pigeons in experiments, all of their naturally occurring “background” lice were eradicated by housing birds in low humidity conditions (< 25% relative ambient humidity) for ≥ 10 weeks. This method kills lice and their eggs, while avoiding residues from insecticides^31^. During experiments, relative humidity in the animal rooms was increased to 35-60%, which provides sufficient humidity for feather lice to extract the moisture they need from the air^32^.

### Measuring louse body size

To measure louse body size, lice were first removed from hosts by anesthetizing them with CO_2_, then ruffling the feathers of the host over a collection tray^33^. Each live louse was photographed by placing it dorsal side up on a glass slide. The lice were harmlessly immobilized by placing a 22 × 22 mm micro cover slip (VWR^®^) directly on the body. Digital photographs were taken at high resolution (uncompressed TIFF 2560 × 1920 pixels) using a DP25 digital camera mounted on an Olympus^®^ SZ-CTV stereoscope linked to a computer running CellSens^®^ image acquisition and analysis software. All of the photos were measured digitally using the open source imaging software ImageJ 1.3. From each image, we measured three aspects of body size: total body length, metathorax width, and head width.

### Experimental evolution of louse body size

To test the influence of host size on louse body size, we infested giant runt pigeons with *C. columbae*. We transferred 800 lice from wild caught feral pigeons to 16 giant runt pigeons and 16 feral pigeon controls (25 lice per bird). At this time (Time 0), we also randomly sampled 800 lice from the source population on wild caught feral pigeons and measured their body size. Pigeons were housed in groups of four in 1.8 × 1.5 × 1.0 m aviaries. Thus, the 32 pigeons used in the experiment were housed in 8 aviaries, each containing four birds of the same type. Housing birds in groups reduced the risk of extinction of experimental lineages of lice on individually housed birds. The experiment ran for 48 mo. *Columbicola columbae* has a mean generation time of 24.4 days^31^; hence, the experiment represents ca. 60 generations of lice.

During the experiment, all pigeons were maintained on a 12-hour light/dark photoperiod and provided *ad libitum* grain, grit, and water. When a bird died during the course of the experiment, lice from the dead bird were transferred to a new parasite-free pigeon of the same type. *Columbicola columbae* can survive for several days on a dead bird, yet cannot leave the bird’s feathers under their own power. Thus, few lice were lost.

Every six months, random samples of lice were removed from pigeons and digitally photographed. To calculate an index of overall body size, we combined measures of total body length, metathorax width, and head width in a principal component analysis (PCA) in JMP v13^34^ (Table S1).

We used linear mixed effects models (LMMs) to explore the relationship between host size and louse size over the course of the experiment. We first quantified experimental changes in overall louse body size (PC1) using an LMM that combined male and female lice from feral and giant runt pigeons. We predicted louse size by modeling the fixed-effects of host type, time (months), louse sex, and all respective interactions, while lineage and aviary were included as random-effects with lineage “nested” within aviary (Table S2). The random effects were included to account for both repeated measures and the structured nature of the data. Three additional LMMs were used to quantify changes in body length, metathorax width, and head width over the course of the experiment (Tables S3-S5). The intercept of each model was set to the value of female lice on feral pigeons at the end of the experiment (48 mo.). All LMMs were fit in R using the “lme4” library package^35,36^. Degrees of freedom and resulting *p*-values were calculated with a Satterthwaite approximation using the “lmerTest” library package^37^.

### Heritability of louse size

Half way through the 48-month study, lice were randomly sampled using the CO_2_ procedure described above. A subsample of adult lice was marked by clipping setae with retinal scissors. Setal clipping is a reliable method that has been used with other species of lice, even under field conditions^38^. Removal of setae does not influence survival, and the setae do not grow back. After clipping, lice from each aviary were combined on a single louse-free common garden feral pigeon. The eight common garden birds were isolated in eight wire mesh cages (30 × 30 × 56 cm). Their preening was impaired to prevent them from removing lice. Preening was impaired using harmless poultry bits, which are C-shaped pieces of plastic inserted between the upper and lower mandibles of a bird’s beak. Bits spring shut in the nostrils to prevent dislodging, but without damaging the tissue. They create a 1-3 mm gap that prevents the forceps-like action of the bill required for efficient preening^39^. Bits have no apparent side effects and they do not impair the ability of birds to feed^40^.

After a period of 48 days, all lice were removed from each of the eight pigeons using CO_2_. At this point in time, most F_1_ offspring had developed to the adult stage, and could be distinguished from members of the parental cohort, which had clipped setae. In contrast, F_2_ lice had not yet developed to the adult stage. Thus, we were able to compare the morphology of parental and F_1_ cohorts of lice from each common garden bird. Parental and F_1_ lice were removed from each bird and digitally photographed and their size measured.

Since *C. columbae* are sexually dimorphic, the body size of F_1_ and parental cohorts were compared in a 2 × 2 matrix: daughters vs. fathers, daughters vs. mothers, sons vs. fathers, and sons vs. mothers. We recovered just one son (F_1_ male) from one of the common garden birds; this common garden bird was excluded from analyses. Hence, son vs father and son vs mother comparisons were restricted to lice from 7 of the 8 common garden birds.

The body size (PC1) of the F_1_ cohort from each common garden pigeon (n = 8-30 lice per common garden bird for a total of 139 lice) was then compared to the body size (PC1) of the parental cohort (n = 11-50 lice per common garden bird for a total of 275 lice). The distribution of body sizes within both parents and offspring were normally distributed. Thus, we used a series of linear regressions to assess the relationship between the mean parental body sizes and the mean offspring body sizes.

### Typical variation in sexual size dimorphism of *C. columbae*

We used walk-in traps to capture 22 feral pigeons in Salt Lake City, Utah. We used the CO_2_ method to recover 262 adult *C. columbae* (roughly equal sex ratio). The 22 pigeons had a mean (± se) of 11.9 (± 1.5) adult lice, reflecting typical adult population sizes at this location^31^. For each population, we subtracted the mean *C. columbae* female length from the mean *C. columbae* male length to generate 22 sexual dimorphism scores (Table S6).

### Influence of sexual size dimorphism on copulation time

We filmed the behavior of arranged pairs of lice that varied in size. We used lice from a group of bitted feral pigeons established for this purpose (but different from the birds used in the common garden experiment). Because bits relax preening-mediated selection for small size, the size of lice on bitted pigeons is similar to, but does not exceed, the size of lice on giant runt pigeons. Each pair of lice was placed in a 9 × 12 mm arena on a detached underwing covert feather. The lice were filmed with an Apple^®^ iPod Touch (5^th^ generation) mounted on an Olympus^®^ SZ-25 stereoscope at 2x magnification for 60min. The videos were watched by two of us (JCA or LIM) and copulation time was recorded. Dimorphism among the pairs of lice was normally distributed. Thus, we used a quadratic regression to examine the relationship between dimorphism and time spent copulating.

### Influence of sexual size dimorphism on egg production

We tracked the egg laying success of 58 arranged pairs of lice that varied in size. Female lice are capable of storing sperm^41^; we therefore bred virgin lice by removing immatures from bitted feral pigeons and reared them to the adult stage on feathers in glass vials in an incubator^18^. Lice were then paired in new glass vials containing feathers. Once the female died, each vial was thoroughly examined for eggs by an observer who was blind to the dimorphism scores (HEC or EJP). Because we scored eggs as present or absent, the egg data had a binomial distribution. Thus, we used an ordinal logistic regression to explore the relationship between dimorphism and likelihood of producing eggs.

### Influence of sexual size dimorphism on reproductive success on live birds

We measured the reproductive success of 36 arranged pairs of lice from giant runt and feral pigeons. We tested whether local adaptation to different sized hosts results in reproductive isolation. We removed immature lice from giant runt and feral pigeons and reared them individually to the adult stage on feathers in glass vials in the incubator^18^. We began with 100 immature lice from each type of host, only half of which survived to adulthood. To avoid removing lice from our experimental evolution lines, we obtained the lice from populations cultured for this purpose on additional giant runt and feral pigeons in our facility.

We infested each of 36 louse-free feral pigeons with a single pair of lice: 12 birds had a male louse from a giant runt and a female louse from a feral pigeon; 12 birds had a female louse from a giant runt and a male louse from a feral pigeon; 12 birds had a male and female louse from a feral pigeon (Table S7). The 36 birds were isolated in 36 cages for 14 days, sufficient time for the lice to breed, but not enough time for offspring to reach the adult stage. After 14 days, the pigeons were sacrificed and the offspring produced by each pair were removed by “body washing” and counted^42^. To test if pairs within the typical range of dimorphism were more likely to have offspring than mismatched pairs, we used a Fisher’s Exact Test.

### Influence of sexual size dimorphism on male-male competition

We filmed the behavior of different sized male lice in the presence of a single female. Lice in this experiment came from bitted feral pigeons. We arranged 30 trios of lice, each consisting of two males and a single female. In each trio, one of the two males showed typical dimorphism relative to the female. The second male was smaller than the typical male, larger than the typical male, or, in the case of controls, was a second typical male (n = 10 for each combination).

Each trio of lice was placed in a 9 × 12 mm arena on a detached underwing covert feather. Behavior of the lice was filmed with an Apple^®^ iPod Touch (5^th^ generation) mounted on an Olympus^®^ SZ-25 stereoscope at 2x magnification for 60min. The videos were watched by two of us (JCA or LIM) and copulation time was recorded. We explored the relationship between size of males relative to females and how much time they spent copulating. This relationship was assessed with a Wilcoxon-Sign Rank Test as the copulation times were not normally distributed and trials were of a paired designed.

## SUPPLEMENTAL INFORMATION

**Table S1.** Principal component analysis (PCA) of louse body size. The PCA is based on the body length, metathorax width, and head width of 3096 lice measured at six-month intervals over the course of the experiment.

|  |  | Principal components |  |  |
| --- | --- | --- | --- | --- |
|  |  | PC1 | PC2 | PC3 |
| Eigenvalues |  | 2.382 | 0.338 | 0.280 |
| % variation |  | 79.4% | 11.3% | 9.3% |
| Eigenvectors | Body length | 0.585 | - 0.126 | - 0.801 |
|  | Metathorax width | 0.571 | 0.765 | 0.297 |
|  | Head width | 0.576 | - 0.631 | 0.519 |
| Loadings | Body length | 0.903 | - 0.073 | - 0.424 |
|  | Metathorax width | 0.882 | 0.445 | 0.157 |
|  | Head width | 0.889 | - 0.367 | 0.275 |

**Table S2:** Linear mixed model (LMM) summary comparing the *overall body size (PC1)* of lice on different size pigeons. This LMM is based on the PC1 scores of 3096 lice measured over the month experiment. Data for lice at Time 0 are a random subsample of lice drawn from the starting population. Data for the rest of the experiment (Time 6 mo. - 48 mo.) are for lice sampled from 32 individual birds (16 feral pigeons, 16 giant runt pigeons) housed in 4 aviaries (4 birds per aviary) for each host treatment.

| *Random effects* | *Variance* | *Std. dev.* |  |  |  |
| --- | --- | --- | --- | --- | --- |
| Lineage (Individual bird) | 0.013 | 0.116 |  |  |  |
| Aviary | 0.027 | 0.165 |  |  |  |
| *Fixed effects* | *Estimate* | *Std. err.* | *df* | *t value* | *Pr(>\|t\|)* |
| Intercept (feral pigeons) | 1.308 | 0.123 | 20.700 | 10.663 | < 0.001* |
| Host (giant runs) | 0.449 | 0.159 | 14.800 | 2.817 | 0.013* |
| Time | 0.006 | 0.002 | 3086.000 | 2.542 | 0.011* |
| Sex (male lice) | -2.437 | 0.119 | 3078.000 | -20.412 | < 0.001* |
| Host x Time | 0.009 | 0.003 | 3086.000 | 3.151 | 0.002* |
| Host x Sex | -0.053 | 0.140 | 3076.000 | -0.377 | 0.706 |
| Time x Sex | 0.001 | 0.003 | 3076.000 | 0.394 | 0.693 |
| Host x Time x Sex | 0.001 | 0.004 | 3073.000 | 0.253 | 0.800 |

\* Indicates significance

**Table S3:** Linear mixed model (LMM) summary comparing the *body length* of lice on different size pigeons. This LMM is based on the length scores of 3098 lice measured over the 48-month experiment. Data for lice at Time 0 are a random subsample of lice drawn from the starting population. Data for the rest of the experiment (Time 6 mo. - 48 mo.) are for lice sampled from 32 individual birds (16 feral pigeons, 16 giant runt pigeons) housed in 4 aviaries (4 birds per aviary) for each host treatment.

| *Random effects* | *Variance* | *Std. dev.* |  |  |  |
| --- | --- | --- | --- | --- | --- |
| Lineage (Individual bird) | 115.500 | 10.750 |  |  |  |
| Aviary | 105.400 | 10.260 |  |  |  |
| *Fixed effects* | *Estimate* | *Std. err.* | *df* | *t value* | *Pr(>\|t\|)* |
| Intercept (feral pigeons) | 2625.854 | 9.056 | 29.200 | 289.953 | < 0.001* |
| Host (giant runts) | 49.380 | 11.548 | 19.400 | 4.276 | < 0.001* |
| Time | -0.568 | 0.186 | 3086.300 | -3.053 | 0.002* |
| Sex (male lice) | -405.548 | 9.623 | 3077.900 | -42.143 | < 0.001* |
| Host x Time | 0.929 | 0.229 | 3088.600 | 4.064 | < 0.001* |
| Host x Sex | -9.383 | 11.311 | 3075.100 | -0.830 | 0.407 |
| Time x Sex | 0.043 | 0.266 | 3075.800 | 0.161 | 0.872 |
| Host x Time x Sex | -0.111 | 0.328 | 3073.200 | -0.338 | 0.735 |

\* Indicates significance

**Table S4:** Linear mixed model (LMM) summary comparing the *metathorax width* of lice on different size pigeons. This LMM is based on the metathorax scores of 3103 lice measured over the 48-month experiment. Data for lice at Time 0 are a random subsample of lice drawn from the starting population. Data for the rest of the experiment (Time 6 mo. - 48 mo.) are for lice sampled from 32 individual birds (16 feral pigeons, 16 giant runt pigeons) housed in 4 aviaries (4 birds per aviary) for each host treatment.

| *Random effects* | *Variance* | *Std. dev.* |  |  |  |
| --- | --- | --- | --- | --- | --- |
| Lineage (individual bird) | 1.003 | 1.001 |  |  |  |
| Aviary | 0.911 | 0.954 |  |  |  |
| *Fixed effects* | *Estimate* | *Std. err.* | *df* | *t value* | *Pr(>\|t\|)* |
| Intercept (feral pigeons) | 297.60 | 1.129 | 70.400 | 263.636 | < 0.001* |
| Host (giant runs) | 4.192 | 1.388 | 41.400 | 3.022 | 0.004* |
| Time | 0.059 | 0.027 | 3067.800 | 2.209 | 0.027* |
| Sex (male lice) | -13.085 | 1.378 | 3088.600 | -9.495 | < 0.001* |
| Host x Time | 0.106 | 0.033 | 3082.900 | 3.231 | 0.001* |
| Host x Sex | -0.223 | 1.620 | 3085.700 | -0.137 | 0.891 |
| Time x Sex | 0.054 | 0.038 | 3086.800 | 1.416 | 0.157 |
| Host x Time x Sex | 0.014 | 0.047 | 3083.900 | 0.304 | 0.762 |

\* Indicates significance

**Table S5:** Linear mixed model (LMM) summary comparing the *head width* of lice on different size pigeons. This LMM is based on the head scores of 3105 lice measured over the 48-month experiment. Data for lice at Time 0 are a random subsample of lice drawn from the starting population. Data for the rest of the experiment (Time 6 mo. - 48 mo.) are for lice sampled from 32 individual birds (16 feral pigeons, 16 giant runt pigeons) housed in 4 aviaries (4 birds per aviary) for each host treatment.

| *Random effects* | *Variance* | *Std. dev.* |  |  |  |
| --- | --- | --- | --- | --- | --- |
| Lineage (individual bird) | 0.587 | 0.767 |  |  |  |
| Aviary | 2.654 | 1.629 |  |  |  |
| *Fixed effects* | *Estimate* | *Std. err.* | *df* | *t value* | *Pr(>\|t\|)* |
| Intercept (feral pigeons) | 288.072 | 1.044 | 13.600 | 275.940 | < 0.001* |
| Host (giant runts) | 1.821 | 1.390 | 10.700 | 1.310 | 0.218 |
| Time | 0.067 | 0.017 | 3091.700 | 4.105 | < 0.001* |
| Sex (male lice) | -10.962 | 0.862 | 3087.800 | -12.712 | < 0.001* |
| Host x Time | 0.023 | 0.020 | 3092.400 | 1.113 | 0.266 |
| Host x Sex | -0.306 | 1.013 | 3085.200 | -0.302 | 0.763 |
| Time x Sex | -0.014 | 0.024 | 3085.500 | -0.577 | 0.564 |
| Host x Time x Sex | 0.010 | 0.029 | 3083.000 | 0.344 | 0.731 |

\* Indicates significance

**Table S6:** Typical variation in body length dimorphism of *C. columbae*. Data are based on the length of 262 adult *C. columbae* removed from 22 wild-caught feral pigeons in Salt Lake City, Utah. All of the lice collected from each bird were used. We subtracted the mean *C. columbae* female length from the mean *C. columbae* male length of lice from each bird to generate 22 sexual dimorphism scores. These data were used to represent the natural range in size dimorphism between male and female *C. columbae* (grey shaded regions in Fig. 4a-c).

| Feral pigeon | Male lice (n) | Female lice (n) | Mean male length (μm) | Mean female length (μm) | Mean body length dimorphism (μm) |
| --- | --- | --- | --- | --- | --- |
| 1 | 5 | 5 | 2015 | 2507 | -492 |
| 2 | 3 | 4 | 2114 | 2572 | -458 |
| 3 | 2 | 1 | 2160 | 2604 | -444 |
| 4 | 4 | 3 | 2158 | 2588 | -430 |
| 5 | 1 | 4 | 2150 | 2573 | -423 |
| 6 | 2 | 8 | 2173 | 2586 | -413 |
| 7 | 2 | 14 | 2162 | 2571 | -409 |
| 8 | 10 | 17 | 2178 | 2555 | -377 |
| 9 | 5 | 2 | 2193 | 2545 | -352 |
| 10 | 2 | 1 | 2227 | 2574 | -347 |
| 11 | 5 | 6 | 2231 | 2566 | -335 |
| 12 | 10 | 4 | 2153 | 2481 | -328 |
| 13 | 10 | 10 | 2217 | 2540 | -323 |
| 14 | 10 | 10 | 2229 | 2543 | -314 |
| 15 | 10 | 5 | 2204 | 2516 | -312 |
| 16 | 2 | 3 | 2227 | 2539 | -312 |
| 17 | 14 | 10 | 2230 | 2539 | -309 |
| 18 | 5 | 5 | 2206 | 2513 | -307 |
| 19 | 3 | 2 | 2227 | 2515 | -288 |
| 20 | 4 | 5 | 2231 | 2514 | -283 |
| 21 | 13 | 6 | 2212 | 2486 | -274 |
| 22 | 10 | 5 | 2242 | 2510 | -268 |

**Table S7:**
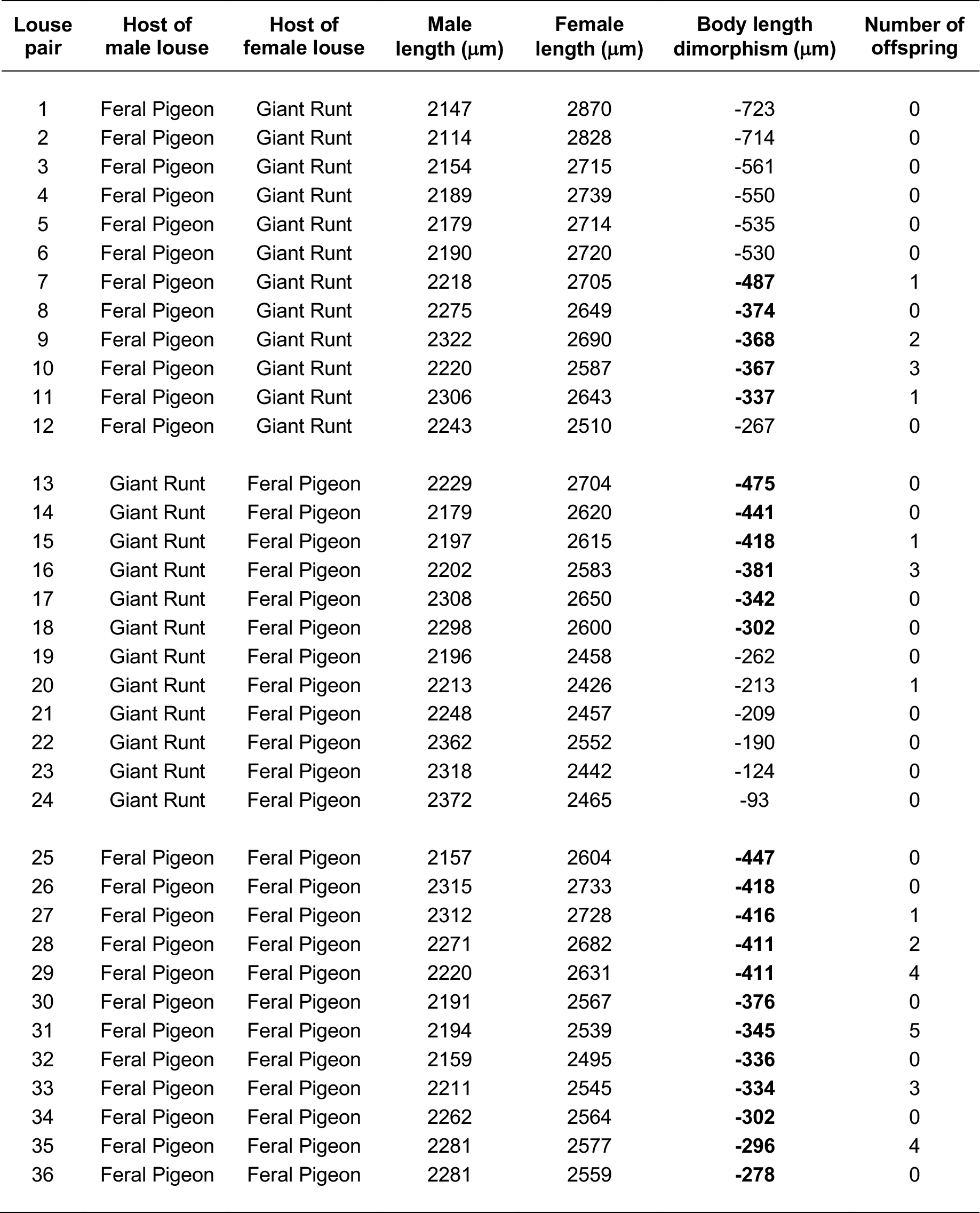
Number of offspring produced by single pairs of lice placed on individually caged feral pigeons (Fig. 4c). Body length dimorphism is the difference in length between the male and female louse of each pair. Boldface values are pairs that fall within the typical range of dimorphism (Fig. 4a-c, Table S6).

### Supplemental Video Legends

**Video S1:**“Typical” sized male and female *C. columbae* copulating. Female on top, male on bottom.

**Video S2:**Relatively large male attempting to copulate. Female on top, male on bottom. The male’s abdomen is too long for sustained copulation.

**Video S3:**Relatively small male attempting to copulate. Female on top, male on bottom. The abdomen of the male is too short for copulation.

**Video S4:**Male-male competition in *C. columbae.* One male displaces another male that is already copulating with a female.

